# Factors Driving Unique Urination Phenotypes of Male and Female 9-week-old C57BL/6J Mice

**DOI:** 10.1101/641167

**Authors:** Hannah Ruetten, Kyle A. Wegner, Helen L. Zhang, Peiqing Wang, Jaskiran Sandhu, Simran Sandhu, Brett Mueller, Zunyi Wang, Jill Macoska, Richard E. Peterson, Dale E. Bjorling, William A. Ricke, Paul C. Marker, Chad M. Vezina

## Abstract

Laboratory mice are used to identify causes of urinary dysfunction including prostate-related mechanisms of Lower Urinary Tract Symptoms (LUTS). Effective use of mice for this purpose requires a clear understanding of molecular, cellular, anatomical, and endocrine contributions to voiding function. Whether the prostate influences baseline voiding function has not been specifically evaluated, in part because most methods that alter prostate mass also change circulating testosterone concentrations. We performed void spot assay and cystometry to establish a multi-parameter “baseline” of voiding function in intact male and female 9-week-old (adult) C57BL/6J mice. We then compared voiding function in intact male mice to that of castrate males, males (and females) treated with the steroid five alpha reductase inhibitor finasteride, or males harboring alleles (*Pbsn4^cre/+^*;*R26R^Dta/+^*) that significantly reduce prostate lobe mass by depleting prostatic luminal epithelial cells. We evaluated aging-related changes in male urinary voiding. We also treated intact male, castrate male, and female mice with exogenous testosterone to determine the influence of androgen on voiding function. The three methods used to reduce prostate mass (castration, finasteride, *Pbsn4^cre/+^*; *R26R^Dta/+^*) changed voiding function from baseline but in a nonuniform manner. Castration feminized some aspects of male urinary physiology (making them more like intact female) while exogenous testosterone masculinized some aspects of female urinary physiology (making them more like intact male). Our results provide evidence that circulating testosterone is responsible in part for baseline sex differences in C57BL/6J mouse voiding function while prostate lobe mass in young, healthy adult mice has a lesser influence.

## 1. INTRODUCTION

Lower urinary tract symptoms (LUTS) are prevalent in adult men and women, negatively affect quality of life, and associate with depression.^1, 2^ It was once believed that LUTS in aging men derive almost exclusively from prostatic enlargement and urethral occlusion.^3^ The rationale was that prostatic volume^4^ and LUTS prevalence and severity^5, 6^ increase with age, and that prostate resection generally alleviates LUTS in aging males.^7^ Recent studies suggest a disease process more complex than previously appreciated. Prostatic volume does not strongly correlate with urodynamic patterns or LUTS severity when measured in the same cohort of men.^5, 8^ Some men with above average prostate volume do not experience clinically significant LUTS while others with below average prostate volumes experience severe LUTS.^9, 10^ It is now becoming clear that male LUTS arise from multiple mechanisms in addition to prostatic enlargement.^11^ There is growing support for the roles of prostatic urethral collagen accumulation,^12^ inflammation,^13^ and smooth muscle hypercontracticility^14, 15^ in progressive LUTS. There is also a growing need for validated model systems to rigorously test these new mechanisms and new pharmacological interventions.

The use of mice to study urinary voiding dysfunction is extensive and widespread. A search of the term “Urology” in PubMed revealed 17,815 publications in 2018: 27 were studies in dog, 575 in rat, and 1003 in mouse. The practice of using male mice for human urinary voiding translational studies has been controversial because not all aspects of mouse and human prostate anatomy are the same. Similarities include anatomical location (base of the bladder) and a narrowing of the urethra in the region where prostatic ducts drain (prostatic urethra). Major differences include prostatic encapsulation and compaction. A portion of the mouse prostate gland lies within a muscular sphincter (rhabdosphinter) but the majority of prostate tissue branches into four bilaterally symmetrical prostate lobes (anterior, dorsal, lateral, and ventral) that are not encapsulated.^16, 17^ The human prostate gland, in contrast, is spherical and has a fibromuscular capsule.^18^ Several studies demonstrate a clear influence of prostate pathologies on mouse urinary voiding behaviors.^19–22^ Influence of the mouse prostate on baseline voiding function has not been specifically evaluated, in part because many methods to alter prostate mass also alter circulating testosterone concentrations.

To address these issues, we used multiple experimental groups and contemporary methods to determine the influence of androgens and prostatic mass on baseline voiding function, lower urinary tract anatomy, and histology. Our companion publication focuses on histology. This report exclusively discusses anatomy and physiology and provides an expansive urinary physiology data set in wild-type C57BL/6J mice that can be mined for hypothesis generation and validation (Tables 2–10). We first evaluated baseline voiding function in nine-week-old male and female C57BL/6J control mice. We then used three different approaches to reduce prostatic mass in male mice: surgical castration, treatment with a steroid 5 alpha reductase inhibitor (finasteride), and a genetic approach to ablate prostate luminal epithelial cells. We also treated intact male, castrate male, and female mice with testosterone to control for the influence of androgen. We identified clear sex differences in baseline urinary function, even though relative bladder mass and volume do not significantly differ between male and female mice. Exogenous testosterone changes several parameters of female mouse urinary function in the direction of intact males (masculinization). Castration feminizes male mouse urinary function, but neither finasteride treatment nor a genetically-induced reduction in prostate mass feminizes male mouse urinary function. We conclude that circulating testosterone is responsible, in part, for sex differences in baseline mouse voiding function while prostatic lobe mass plays a lesser role.

## 2. MATERIALS AND METHOD

### 2.1 Mice

All experiments were conducted under a protocol approved by the University of Wisconsin Animal Care and Use Committee and in accordance with the National Institutes of Health Guide for the Care and Use of Laboratory Animals. Mice were housed in Udel^®^ Polysulfone microisolator cages on racks or in Innocage^®^ disposable mouse cages on an Innorack^®^; room lighting was maintained on 12 hour light and dark cycles; room temperature was maintained at 20.5 ± 5 C; humidity was 30–70%. Mice were fed 8604 Teklad Rodent Diet (Harlan Laboratories, Madison WI) and feed and water were available *ad libitum*. Cages contained corn cob bedding. All endpoint measurements were collected in nine-week-old mice unless specified otherwise.

All mice used in this study were purchased from Jackson Laboratories (Bar Habor, ME) and included C57BL/6J (Stock #000664), Tg(Pbsn-*cre*)4Prb/J (Pbsn4*cre*, stock number 026662) bred onto the C57BL/6J background for four to five generations,^23^ B6.Cg-Gt(ROSA)26Sor^tm14(CAG-tdTomato)Hze^/J (R26R-Tdtomato, Jax stock #007914),^24^ and Gt(ROSA)26Sortm1(DTA)Jpmb/J (R26R-Dta, Jax stock #006331)^25^. The genotype of mice with depleted prostatic epithelial cells was Pbsn4^cre/+^; R26R^Dta/+^ and their respective littermate controls were Pbsn4^cre/+^; R26R^TdTomato/+^.

### 2.2 Castration/Sham Castration

Castration was performed at six weeks of age. Mice were anesthetized with isoflurane and given ketoprofen (0.5 mg/kg sc) as an analgesic. A midline incision was made in the scrotum, and the testes were either removed (castrate) or examined (sham controls). The scrotum was closed using a simple interrupted pattern.

### 2.3 Testosterone Capsule Preparation and Implantation

Capsules were prepared as previously described.^26^ Silastic tubing (Dow Corning, Cat. # 508-008, Silastic Laboratory Tubing, 1.57 mm inside diameter X 3.18 mm outside diameter) was cut to 16 mm. The wooden stick of cotton tipped applicators (Fisher Scientific Cat. #23-400-100) was cut into 5 mm pieces and inserted 3 mm into the tubing to plug the ends. 10 mm of the sham capsule was left empty. 10 mm of the testosterone capsule was filled with 4-androsten-17beta-ol-3-one (Testosterone, T, ≥99% pure, Steraloids Inc. Cat. # A6950-000). Silastic capsules were sealed with silastic medical adhesive, type A (Dow Corning, purchased from Factor II Inc, [Product No. A-100]). Testosterone filled silastic capsules have been shown to effectively increase testosterone in C57BL/6J mice when implanted as described.^27^

Mice were anesthetized with isoflurane for silastic capsule implantation and given ketoprofen (0.5 mg/kg sc). An incision was made on the caudal aspect of the back just to the right of midline. Capsules were inserted parallel to the spine and the incision was closed with wound clips.

### 2.4 Finasteride Treatment

Finasteride (Alfa Aesar, J63454, Ward Hill, MA) was dissolved in 100% EtOH and diluted in corn oil to make a 10% EtOH/ 90% corn oil dosing solution. The solution was stored at 4°C for the duration of the experiment. Mice were given finasteride (50 µL of a 40 µg/µL solution) daily via oral gavage.

### 2.5 Void Spot Assay

We followed the recommended guidelines of reporting VSA data.^28–30^ VSA was performed in the vivarium where mice were housed one day prior to cystometry and euthanasia. Whatman grade 540 (Fisher Scientific no. 057163-W) filter papers (27 × 16 cm) were placed in the bottom of Udel^®^ Polysulfone microisolator cages. Mice were placed in the cage (singly housed) with food *ad libitum* but no water for four hours starting from 8-11 AM GMT. VSA was performed once a week starting at six weeks of age, allowing for three acclimation sessions prior to the session at nine weeks of age which was used for analysis. Filter papers were dried and imaged with an Autochemi AC1 Darkroom ultraviolet imaging cabinet (UVP, Upland, CA) equipped with an Auto Chemi Zoom lens 2UV and an epi-illuminator. Image capture settings were adjusted using UVP VisonWorksLS image acquisition software. Images were captured using an Ethidium Bromide filter set (570-640 nm) and 365 nm epi-illumination. Void Whizzard was downloaded from http://imagej.net/Void_Whizzard and run according to the user guide.^30^ Analyzed parameters included: Total Spot Count, Total Void Area (cm^2^), % area in center of paper, % area in corners of paper, and mass distribution of spots (0-0.1, 0.1-0.25, 0.25-0.5, 0.5-1, 1-2, 2-3, 3-4, 4+ cm).

### 2.6 Anesthetized Cystometry

Cystometry was performed with minimal alterations to previously published protocols.^31, 32^ Mice were anesthetized with urethane (1.43 g/kg sc). Thirty minutes after urethane dosing, an incision was made in the ventral abdomen to expose the bladder. Bladder length and diameter were measured for volume calculation. A purse-string suture was placed in the bladder dome. Polyethylene cystostomy tubing (PE50, outer diameter 0.58mm, inner diameter 0.28mm) was inserted into the bladder through the center of the suture and purse-string secured to hold the tubing in place with 2-3 mm of tubing within the bladder. The abdominal wall and skin were closed separately in a simple interrupted pattern. The exterior tubing was secured to the ventral abdominal skin with two simple interrupted sutures. Mice were placed on a heat pad for one hour after the procedure.

The exposed tube was connected to a three-way stopcock, and the other two arms of the stopcock were connected to an infusion pump (Harvard Apparatus, Holliston, MA) and pressure transducer (Memscap AS, Norway). Intravesical pressure was recorded continuously using a PowerLab data collection system (ADI Instruments, Colorado Springs, CO). Room-temperature sterile saline (0.9%) was infused into the bladder at a rate of 0.8 mL per hour.

Mice were placed in lateral recumbency above a force transducer (Model FT03, grass Instruments) with a 3D printed urine collection funnel. The force transducer was calibrated with known volumes of saline to create a pressure-volume conversion. The mass of voided urine was recorded continuously using PowerLab.

At least one hour of voiding activity was recorded. Three to five consecutive voids, occurring after stabilization of micturition cycles, were used for analyses. Multiple parameters were measured including: Void Duration, Intervoid Interval, Baseline Pressure, Normalized Threshold Pressure, Normalized Peak Void Pressure, Number of Non-Voiding Contractions, Voided Volume, Compliance, Volume Flow Rate, Mass Based Flow Rate, and Efficiency (Calculations used for analysis of cystometric tracings are described in Figure 1, Table 1).

**Figure 1.**
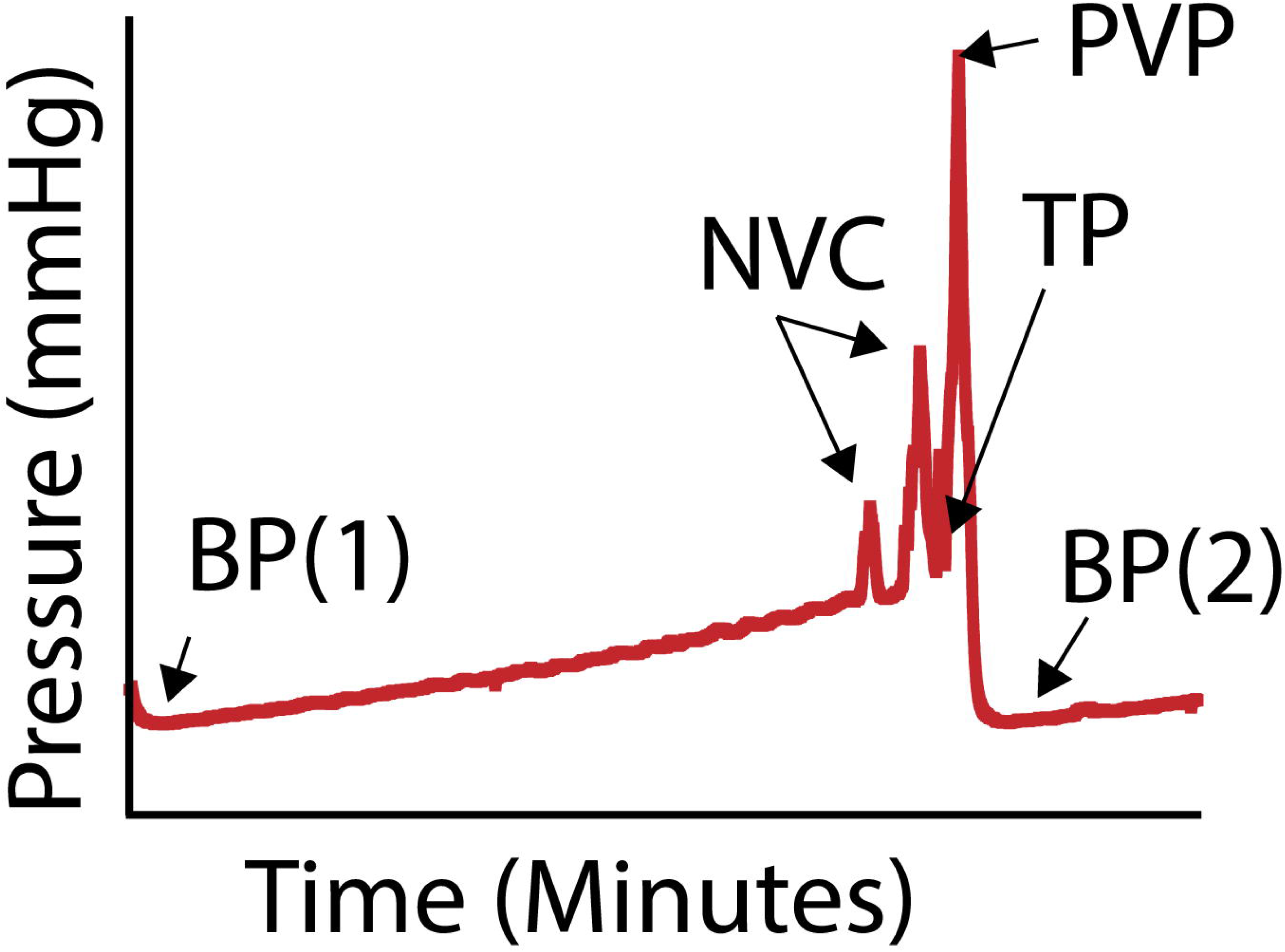
A cystometry trace with annotated features that were used for calculating the cystometry endpoints in Table 1. BP 1 & 2 are baseline pressure, the lowest bladder pressure before bladder filling (BP(1))/ immediately after voiding is complete (BP(2)). NVC is a non-voiding contraction, a bladder pressure spike occurring without urination. TP is the threshold pressure, the pressure when voiding initiates. PVP is the peak void pressure, the maximal bladder pressure achieved during voiding.

**Table 1.** Formulas for Calculated Cystometry Parameters.

| **CMG Parameter** | **Abbreviation** | **Units** | **Description** |
| --- | --- | --- | --- |
| Baseline Pressure | BP | *mmHg* | *lowest pressure of trace immediately after void; also defines starting point of new void (see figure 1, BP)* |
| Number of Non-Voiding Contractions | nNVC | *(no units)* | *Number of spikes in pressure prior to voiding that did not result in urine excretion (see figure 1, NVC)* |
| Threshold Pressure | TP | *mmHg* | *Pressure at start of urination (see figure 1, TP)* |
| Peak Void Pressure | PVP | *mmHg* | *Greatest pressure achieved during void(see figure 1, PVP)* |
| Normalized Peak Void Pressure | NPVP | *mmHg* | $PVP - BP$ |
| Normalized Threshold Pressure | NTP | *mmHg* | $TP - BP$ |
| Void Duration | VD | *min* | $(\text{time at BP of void}_{(n+1)}) - (\text{time at TP of void}_n)$ |
| Voided Volume | VV | *mL* | $(\text{mass of urine trace after void}) - (\text{mass of urine trace prior to void})$ |
| Intervoid Interval | IVI | *min* | $(\text{time at PVP of void}_{(n+1)}) - (\text{time at PVP of void}_n)$ |
| Infusion Rate | IR | *mL/min* | *rate at which the pump infuses saline* |
| Infusion Duration | ID | *min* | $TP - BP$ |
| Infused Volume | IV | *mL* | $IR \times ID$ |
| Efficiency | E | *%* | $\frac{VV}{IV}$ |
| Compliance | C | *mL/mmHg* | $\frac{IV}{(TP - BP)}$ |
| Volume Flow Rate | VFR | *mL/min* | $\frac{VV}{VD}$ |
| Mass Based Flow Rate | MBFR | *mg/min* | *slope of urine mass trace during active voiding* |

### 2.7 Statistical Analysis

Statistical analyses were performed with Graph Pad Prism 8.0.2 (Graphpad Software, La Jolla, California). Shapiro-Wilk test was used to test for normality and transformation was applied to normalize data when possible. The F-test was used to test for homogeneity of variance for pairwise comparisons. Welsh’s correction was applied when variances was unequal. When variance was equal, comparisons between two groups were made using Student’s t-test. The Mann Whitney test was applied when data could not be normalized through transformation. Bartlett’s test was used to test for homogeneity of variance for multiple comparisons. Welsh’s ANOVA was applied when variance was unequal followed by Tamhane’s T2 multiple comparisons test. When variance was equal, comparisons between groups were made using ordinary one-way ANOVA followed by Sidak’s multiple comparisons test. If data could not be normalized through transformation, the Kruskal-Wallis test was applied with Dunn’s multiple comparisons test. A p < 0.05 was considered statistically significant. We have also noted changes that approach significance p < 0.10 when they are consistent with our hypothesis. These changes are marked with “^Δ^” in tables and referred to in text as “trends” accompanied by the appropriate p-value. All numerical data are presented as mean +/- standard error of the mean (SEM).

## 3. RESULTS

### 3.1 Male and Female Mouse Baseline Urinary Voiding Physiology Characteristics

We evaluated several voiding parameters in male and female mice to determine the impact of sex on urinary voiding physiology (Figure 2-A, Figure 4-A, and Table 2). Body mass is greater in males than females, consistent with previous findings in C57BL/6J mice.^31^ Relative bladder mass and volumes were determined by normalizing to body mass. Male and female relative bladder weight and volume do not significantly differ, consistent with previous studies.^33^

**Figure 2.**
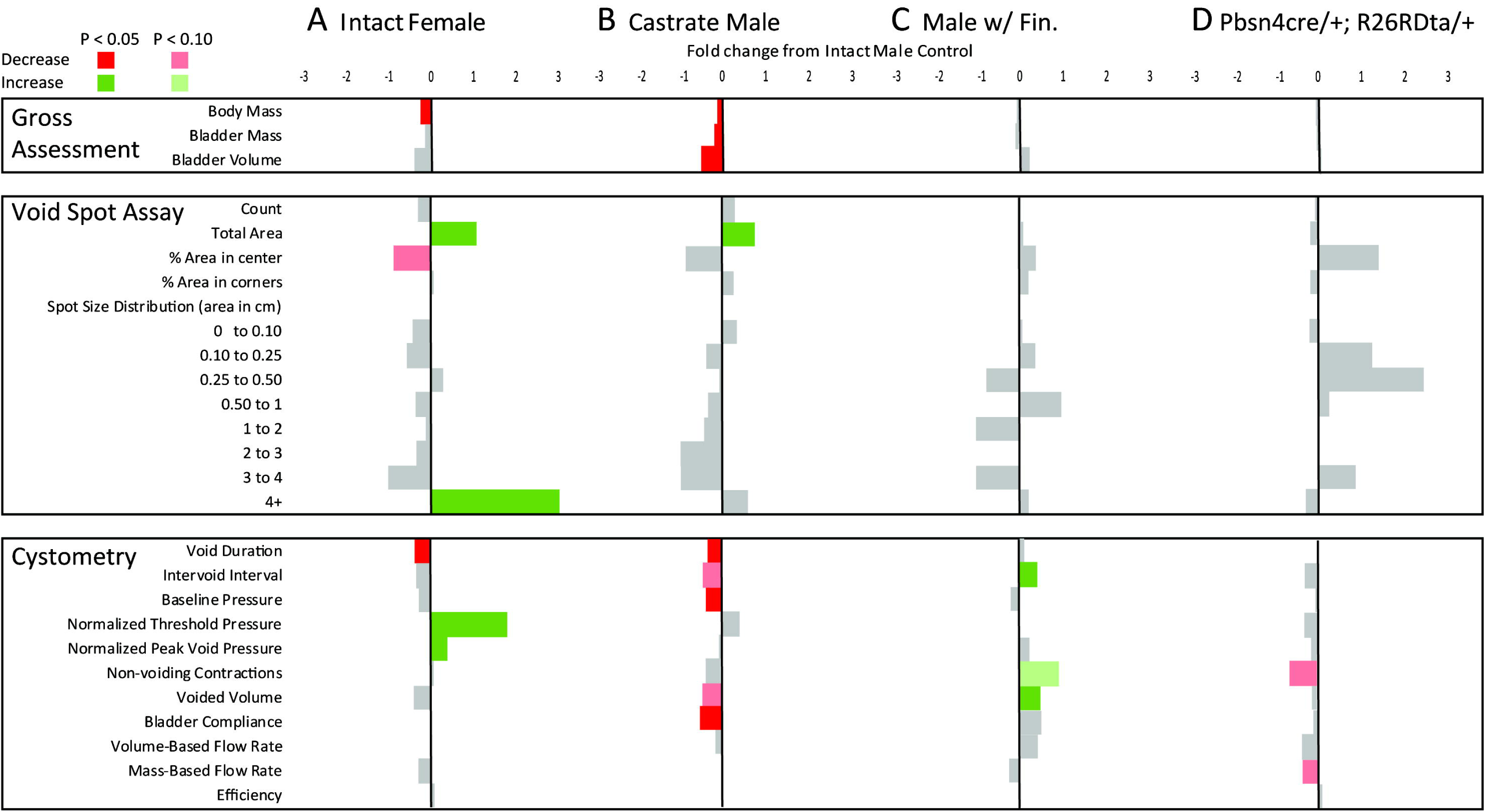
Summary statistics for anatomical and physiological differences across experimental groups evaluated in this study including intact male and female mice, castrate male mice (Castrate Male) mice treated with the steroid 5 alpha reductase inhibitor finasteride (Males w/ Fin), or males genetically engineered to produce diphtheria toxin in luminal epithelial cells, resulting in their depletion (Pbsn4cre/+;R26RDta/+). All results are mean fold differences relative to intact male control. Differences that are significantly higher than intact males are shown in green and those that are lower than intact males are in red. Endpoints that do not significantly differ from intact males are in gray. The numerical fold difference relative to intact males is indicated by the bar size according to the legend at the top of the figure. Female urinary physiology is different from male at baseline. Castration causes some changes consistent with feminization of urinary function. Three different methods of prostate mass reduction held no consistent effects on urinary physiology.

**Table 2.** Male and Female C57BL/6J Mouse Baseline Urinary Voiding Characteristics.

| **Gross Assessment** | | Values are: Average ± SEM | |
| --- | --- | --- | --- |
|  | Intact Male<br>(n = 13) | Intact Female<br>(n = 7) | p-value |
| **Body Mass (g)** | **24.45 ± 0.49** | **18.13 ± 0.40** | **<0.0001** |
| Bladder |  |  |  |
| Mass (% Body Mass) | 0.15 ± 0.01 (n=6) | 0.13 ± 0.01 | 0.1639 |
| Volume (mm<sup>3</sup>/g Body Mass) | 39.86 ± 7.46 | 22.72 ± 5.41 | 0.1366 |
| **Void Spot Assay** |  |  |  |
|  | Intact Male<br>(n = 9) | Intact Female<br>(n = 7) | p-value |
| Count | 32.11 ± 12.67 | 22.86 ± 1.16 | 0.4890 |
| **Total Area (cm<sup>2</sup>)** | **11.49 ± 3.17** | **23.84 ± 2.13** | **0.0089** |
| Percent area in center | 15.78 ± 9.97 | 2.48 ± 1.48 | 0.0524<sup>Δ</sup> |
| Percent area in corners | 61.14 ± 13.26 | 65.12 ± 9.69 | 0.8220 |
| Spots of a certain size (area) |  |  |  |
| 0 - 0.1 cm<sup>2</sup> | 28.11 ± 11.30 | 16.29 ± 0.68 | 0.9740 |
| 0.1 - 0.25 cm<sup>2</sup> | 1.00 ± 0.47 | 1.57 ± 0.43 | 0.2940 |
| 0.25 - 0.5 cm<sup>2</sup> | 0.67 ± 0.33 | 0.86 ± 0.14 | 0.2550 |
| 0.5 - 1 cm<sup>2</sup> | 0.67 ± 0.37 | 0.43 ± 0.20 | >0.9999 |
| 1 - 2 cm<sup>2</sup> | 0.33 ± 0.33 | 0.29 ± 0.18 | 0.5500 |
| 2 - 3 cm<sup>2</sup> | 0.44 ± 0.34 | 0.29 ± 0.29 | 0.8500 |
| 3 - 4 cm<sup>2</sup> | 0.11 ± 0.11 | 0 ± 0 | >0.9999 |
| **4+ cm<sup>2</sup>** | **0.78 ± 0.22** | **3.14 ± 0.26** | **<0.0010** |
| **Cystometry** |  |  |  |
|  | Intact Male<br>(n = 6) | Intact Female<br>(n = 7) | p-value |
| **Void Duration** | **0.64 ± 0.08** | **0.41 ± 0.02** | **0.0252** |
| Intervoid Interval | 5.10 ± 1.18 | 3.50 ± 0.46 | 0.2430 |
| Baseline Pressure | 3.61 ± 0.31 | 2.68 ± 0.81 | 0.2800 |
| **Normalized Threshold Pressure** | **3.35 ± 0.32** | **9.36 ± 1.16** | **<0.0001** |
| **Normalized Peak Void Pressure** | **18.37 ± 2.06** | **25.35 ± 1.11** | **0.0160** |
| Non-voiding contractions | 2.26 ± 0.72 | 2.4 ± 0.61 | 0.8840 |
| Voided Volume ( x 10<sup>-2</sup>) | 7.60 ± 1.74 | 4.72 ± 0.51 | 0.1660 |
| **Compliance ( x 10<sup>-2</sup>)** | **1.92 ± 0.49** | **0.52 ± 0.10** | **0.0025** |
| Volume Flow Rate ( x 10<sup>-3</sup>) | 1.98 ± 0.30 | 1.96 ± 0.21 | 0.9390 |
| Mass Based Flow Rate | 23.70 ± 4.09 | 16.80 ± 3.97 | 0.2530 |
| Efficiency (%) | 96.40 ± 3.98 | 104.20 ± 4.24 | 0.2080 |

We used void spot assay (VSA) as a first approach to evaluate voiding behaviors. The VSA procedure has been refined considerably in recent years. Methodology improvements minimize experimental bias, new software enables rigorous and unbiased assessment, ^28, 30^ and 12 endpoint measurements collected in this study confer a more robust and multidimensional perspective than the five or less measurements collected in prior comparisons of male and female C57BL/6J mice.^31, 33^ We identified several differences between control male and female mice. Female mice deposit more total urine (measured by area, cm^2^) in a four-hour period than males. Our findings differ from that of a previous study involving older mice and using a method of VSA analysis that excluded urine spots <0.66 cm^2^.^31^ It is worth noting that we recently showed that small void spots are not caused by mice tracking urine from deposited voids, a rationale for their previous exclusion.^30^

Previous studies excluding small and large void spots, or spots in cage corners, did not observe male-female differences in voiding patterns.^31, 33^ When we evaluated the spatial distribution of voids (center, corners, and in-between), we found that female mice trend toward depositing less urine in the center of the cage (p = 0.0524). We also evaluated the categorical distribution of VSA spots (0-0.1, 0.1-0.25, 0.25-0.5, 0.5-1, 1-2, 2-3, 3-4, and 4+ cm^2^) and found that female mice deposit a greater proportion of large voids (4+ cm^2^) than males.

We next used anesthetized cystometry (CMG) to evaluate voiding function. This method has also evolved in recent years. New practices include: 1) A novel method of urethane delivery (s.c.) that minimizes spontaneous body movements and their associated influence on intravesicular pressures, 2) computer generated traces that improve accuracy of measured trace characteristics, and 3) simultaneous collection of intravesical pressure and voided urine mass allowing for calculation of additional void characteristics. We now routinely collect 11 CMG endpoint measurements compared to the four to five collected in previous studies. We found that peak void pressures are higher in females than males, consistent with a previous study.^31^ Also consistent with previous studies, we did not observe sex differences in intervoid interval, number of non-voiding contractions, and voided volume.^31, 33^ Novel findings from this study are that female mice have higher threshold pressures, shorter void durations, and less bladder compliance than males.

The mouse urinary voiding behaviors we report in this study are specific to C57BL/6J mice and may be different across the multitude of mouse strains, with mice of difference age, as well as health or disease status. Sex differences and mouse urinary phenotype were previously reported to be strain specific.^31^ Age and disease state in a genetically, surgically, or otherwise altered mice, could also impact sex differences in mice. We recently documented an aging-related voiding dysfunction between 2-month and 24-month-old male C57BL/6J mice.^19^ Here, we honed in on urinary function in young adult (1.5 – 3.5-month-old) mice. We performed VSA on 6, 7, 8, 9, 10, 12, and 14-week old male C57BL/6J mice and all measured endpoints changed with age (Figure 3, Table 3).

**Figure 3.**
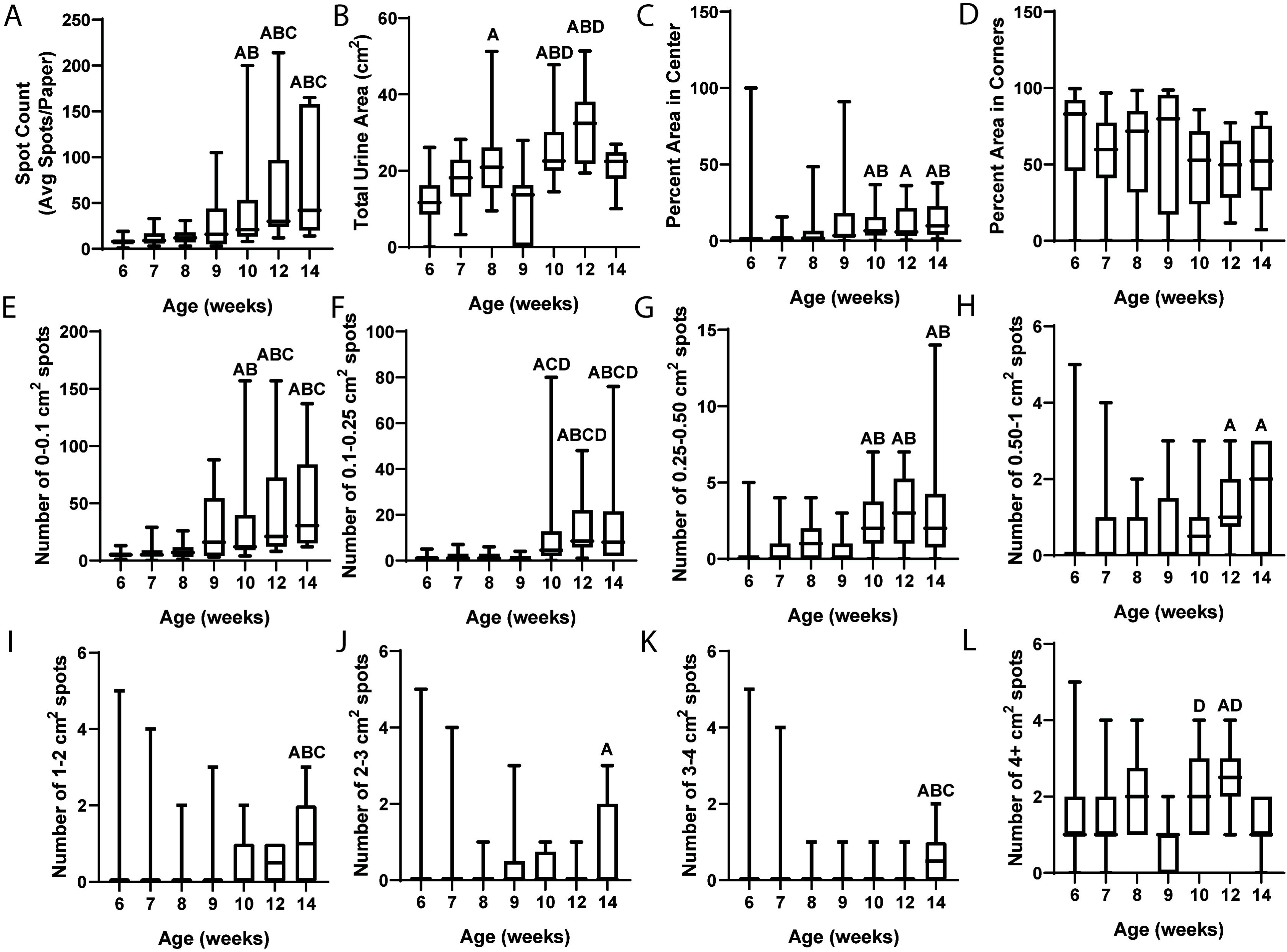
Aging (from 6-to-14-weeks-old) alters void spot assay parameters in male C57BL/6J mice. Thirty mice underwent void spot assay (VSA) at 6 and 7 weeks, twenty at 8 and 10 weeks, and ten at 9, 12, and 14 weeks of age. The relationship between age and each VSA parameter is shown as box-and-whisker plots. Age groups were compared and A- designates a significant difference from 6-week-old mice, B- from 7-week-old mice, C- from 8-week-old mice, and D- from 9 week-old-mice. Six to nine-week-old mice, and ten to fourteen-week-old mice are statistically similar in void spot assay characteristics. (A) Spot count, (B) total urine area, (C) percent area in center, and (E-L) all spot sizes are altered with age.

**Table 3.** Impact of Age on VSA Characteristics.

| Void Spot Assay | Values are: Average ± SEM | | | | | | |
| --- | --- | --- | --- | --- | --- | --- | --- |
|  | 6-week-old | 7-week-old | 8-week-old | 9-week-old | 10-week-old | 12-week-old | 14-week-old |
| Count | 8.33 ± 0.88 | 11.97 ± 1.50 | 13.20 ± 1.66 | 32.10 ± 12.70 | 53.45 ± 13.66<sup>A,B</sup> | 65.90 ± 20.43<sup>A,B,C</sup> | 75.10 ± 19.38<sup>A,B,C</sup> |
| Total Area (cm<sup>2</sup>) | 12.68 ± 1.07 | 17.34 ± 1.22 | 21.89 ± 2.28<sup>A</sup> | 11.50 ± 3.17 | 25.91 ± 1.88<sup>A,B,D</sup> | 31.69 ± 3.08<sup>A,B,D</sup> | 21.01 ± 1.70 |
| Percent area in center | 5.26 ± 3.32 | 2.41 ± 0.69 | 6.31 ± 2.57 | 15.80 ± 9.97 | 10.74 ± 2.51<sup>A,B</sup> | 11.38 ± 3.95<sup>A</sup> | 14.08 ± 4.08<sup>A,B</sup> |
| Percent area in corners | 69.67 ± 5.94 | 59.22 ± 4.30 | 57.87 ± 7.45 | 61.10 ± 13.30 | 48.17 ± 6.24 | 47.64 ± 7.06 | 51.56 ± 8.19 |
| Spots of a certain size (area) |  |  |  |  |  |  |  |
| 0 - 0.1 cm<sup>2</sup> | 5.37 ± 0.65 | 7.57 ± 1.17 | 8.10 ± 1.29 | 28.10 ± 11.30 | 35.45 ± 9.90<sup>A,B</sup> | 44.20 ± 15.56<sup>A,B,C</sup> | 50.20 ± 14.15<sup>A,B,C</sup> |
| 0.1 - 0.25 cm<sup>2</sup> | 1.10 ± 0.23 | 1.80 ± 0.35 | 1.55 ± 0.39 | 1.00 ± 0.47 | 11.95 ± 4.40<sup>A,C,D</sup> | 14.30 ± 4.47<sup>A,B,C,D</sup> | 16.20 ± 7.13<sup>A,B,C,D</sup> |
| 0.25 - 0.5 cm<sup>2</sup> | 0.23 ± 0.11 | 0.37 ± 0.16 | 1.00 ± 0.26 | 0.67 ± 0.33 | 2.25 ± 0.38<sup>A,B</sup> | 3.00 ± 0.75<sup>A,B</sup> | 3.30 ± 1.29<sup>A,B</sup> |
| 0.5 - 1 cm<sup>2</sup> | 0.20 ± 0.07 | 0.47 ± 0.16 | 0.45 ± 0.14 | 0.67 ± 0.37 | 0.75 ± 0.20 | 1.20 ± 0.29<sup>A</sup> | 1.60 ± 0.40<sup>A</sup> |
| 1 - 2 cm<sup>2</sup> | 0.07 ± 0.05 | 0.23 ± 0.16 | 0.15 ± 0.11 | 0.33 ± 0.33 | 0.50 ± 0.17 | 0.50 ± 0.17 | 1.10 ± 0.35<sup>A,B,C</sup> |
| 2 - 3 cm<sup>2</sup> | 0 ± 0 | 0.07 ± 0.05 | 0.05 ± 0.05 | 0.44 ± 0.34 | 0.25 ± 0.10 | 0.10 ± 0.10 | 0.80 ± 0.36<sup>A</sup> |
| 3 - 4 cm<sup>2</sup> | 0.03 ± 0.03 | 0.07 ± 0.05 | 0.05 ± 0.05 | 0.11 ± 0.11 | 0.15 ± 0.08 | 0.10 ± 0.10 | 0.60 ± 0.22<sup>A,B,C</sup> |
| 4+ cm<sup>2</sup> | 1.33 ± 0.12 | 1.40 ± 0.14 | 1.85 ± 0.21 | 0.78 ± 0.22 | 2.15 ± 0.22<sup>D</sup> | 2.50 ± 0.27<sup>A,D</sup> | 1.30 ± 0.21 |

<sup>A</sup> -Significant difference from 6-week-old <sup>B</sup> -Significant difference from 7-week-old <sup>C</sup> -Significant difference from 8-week-old <sup>D</sup> -Significant difference from 9-week-old

### 3.2 Prostate Mass Reduction Minimally Impacts Baseline Male Mouse Urinary Physiology

We hypothesized that the prostate may contribute to sex differences in C57BL/6J mouse voiding behaviors. We used three strategies to reduce prostate mass and evaluate the resulting impact on voiding function: Castration, 5 alpha reductase inhibitor (finasteride) treatment, and genetic prostatic luminal cell ablation. We specifically tested whether our three methods reduce prostate mass as expected, cause a consistent directional change in VSA and/or CMG voiding characteristics, and whether the directional change is consistent with “feminization” of urinary physiology (i.e. same directional change as female compared to male).

The influence of castration on male voiding function is summarized in Figure 2-B and Table 4. Castration, more than any other method to reduce prostate lobe mass, causes the greatest magnitude of prostate lobe mass reduction, but also reduces body, bladder, seminal vesicle mass, and bladder volume as reported previously in Swiss mice.^34, 35^ Castrate and intact male mice did not overwhelmingly differ in VSA-measured voiding function, but there were some differences consistent with “feminization” of male urinary physiology. Castrate male mice, like female mice, deposit more urine in a four-hour monitoring period compared to intact males. This is consistent with a previous finding of increased urine mass/time in castrate Swiss mice.^35^ Castrate mice are similar to intact males in the percent of voids deposited in the center of the cage and the number of 4+ cm^2^ spots. Castrate males and intact females have a shorter void duration and a lower bladder compliance than intact males. However, castrate males have a lower baseline bladder pressure than intact control males, an endpoint that distinguishes them from females. Additionally, castrate males trend toward a shorter intervoid interval and lower voided volume than intact males (p = 0.0968 and p = 0.0977 respectively).

**Table 4.** Impact of Castration on Male Urinary Physiology.

| **Gross Assessment** | | Values are: Average ± SEM | |
| --- | --- | --- | --- |
|  | Intact Male<br>(n = 13) | Castrate Male<br>(n = 14) | p-value |
| **Body Mass (g)** | **24.45 ± 0.49** | **21.51 ± 0.41** | **<0.0001** |
| **Bladder** |  |  |  |
| Mass (% Body Mass) | 0.15 ± 0.01 (n=6) | 0.12 ± 0.00 (n=6) | 0.0190 |
| Volume (mm<sup>3</sup>/g Body Mass) | 39.86 ± 7.46 | 18.04 ± 3.13 | 0.0041 |
| **Prostate Mass (% Body Mass)** | **0.20 ± 0.01** | **0.03 ± 0.00** | **<0.0001** |
| Anterior Mass (% Body Mass) | 0.09 ± 0.00 | 0.02 ± 0.00 | <0.0001 |
| Ventral Mass (% Body Mass) | 0.04 ± 0.00 | 0.01 ± 0.00 | <0.0001 |
| Dorsal Mass (% Body Mass) | 0.04 ± 0.00 | 0.01 ± 0.00 | <0.0001 |
| Lateral Mass (% Body Mass) | 0.01 ± 0.00 | 0.00 ± 0.00 | 0.0003 |
| **Seminal Vesicle Mass (% Body Mass)** | **0.62 ± 0.04** | **0.04 ± 0.00** | **<0.0001** |
| **Void Spot Assay** |  |  |  |
|  | Intact Male<br>(n = 9) | Castrate Male<br>(n = 11) | p-value |
| Count | 32.10 ± 12.70 | 42.00 ± 6.10 | 0.1025 |
| **Total Area (cm<sup>2</sup>)** | **11.50 ± 3.17** | **20.50 ± 2.31** | **0.0301** |
| Percent area in center | 15.80 ± 9.97 | 2.34 ± 0.666 | 0.1567 |
| Percent area in corners | 61.10 ± 13.30 | 77.40 ± 3.96 | 0.2177 |
| Spots of a certain size (area) |  |  |  |
| 0 - 0.1 cm<sup>2</sup> | 28.10 ± 11.30 | 38.10 ± 5.84 | 0.1472 |
| 0.1 - 0.25 cm<sup>2</sup> | 1.00 ± 0.47 | 1.36 ± 0.43 | 0.5651 |
| 0.25 - 0.5 cm<sup>2</sup> | 0.67 ± 0.33 | 0.64 ± 0.15 | 0.6534 |
| 0.5 - 1 cm<sup>2</sup> | 0.67 ± 0.37 | 0.46 ± 0.21 | 0.9591 |
| 1 - 2 cm<sup>2</sup> | 0.33 ± 0.33 | 0.18 ± 0.12 | >0.9999 |
| 2 - 3 cm<sup>2</sup> | 0.44 ± 0.34 | 0 ± 0 | 0.1895 |
| 3 - 4 cm<sup>2</sup> | 0.11 ± 0.11 | 0 ± 0 | 0.4500 |
| 4+ cm<sup>2</sup> | 0.78 ± 0.22 | 1.27 ± 0.14 | 0.1328 |
| **Cystometry** |  |  |  |
|  | Intact Male<br>(n = 6-7) | Castrate Male<br>(n = 7-8) | p-value |
| **Void Duration** | **0.64 ± 0.08** | **0.42 ± 0.03** | **0.0311** |
| Intervoid Interval | 5.10 ± 1.18 | 2.77 ± 0.12 | 0.0968<sup>Δ</sup> |
| **Baseline Pressure** | **3.61 ± 0.31** | **2.21 ± 0.26** | **0.0040** |
| Normalized Threshold Pressure | 3.35 ± 0.33 | 4.75 ± 0.73 | 0.1087 |
| Normalized Peak Void Pressure | 18.40 ± 2.06 | 17.20 ± 0.75 | 0.6145 |
| Non-voiding contractions | 2.26 ± 0.72 | 1.46 ± 0.25 | 0.3254 |
| Voided Volume (x 10<sup>-2</sup>) | 7.60 ± 1.74 | 4.05 ± 0.11 | 0.0977<sup>Δ</sup> |
| **Compliance (x 10<sup>-2</sup>)** | **1.92 ± 0.49** | **0.89 ± 0.20** | **0.0359** |
| Volume Flow Rate (x 10<sup>-3</sup>) | 1.98 ± 0.30 | 1.69 ± 0.16 | 0.3871 |
| Mass Based Flow Rate | 23.70 ± 4.09 | 23.4 ± 3.17 | 0.9452 |
| Efficiency (%) | 96.40 ± 3.98 | 96.8 ± 3.306 | 0.9381 |

We next reduced prostate mass by treating mice for two weeks with finasteride (100 mg/kg BID via oral gavage) to block conversion of testosterone to the more potent ligand, dihydrotestosterone, and reduce androgen concentration in a non-surgical manner. This dosing paradigm reflects that of a previous study in rats, which reported 24h of sustained serum finasteride concentrations following a single 100 mg/kg oral dose and a prostate mass reduction two weeks after treatment.^36^ Results are summarized in Figure 2-C and Table 5. Males treated with finasteride (Males w/ Fin) have significantly smaller seminal vesicles and smaller anterior, ventral, and lateral prostates than oil-treated control males. Dorsal prostate mass does not differ between groups. VSA-measured voiding function in males w/ Fin does not differ from control males. CMG-measured endpoints differed between groups (males w/ Fin had a significantly larger intervoid interval and voided volume, and trended (p=0.0755) toward more non-voiding contractions than control males). The finasteride-mediated directional changes in voiding function were different than those caused by castration. We treated female mice (which have a very low baseline level of testosterone) with finasteride and performed VSA and CMG to determine the impact of finasteride on female urinary function. Finasteride causes some changes in VSA-measured voiding function in females but no changes in CMG-measured function (Table 6).

**Table 5.** Impact of Finasteride on Male Urinary Physiology.

| **Gross Assessment** | | Values are: Average ± SEM | |
| --- | --- | --- | --- |
|  | **Males w/ Oil**<br>**(n = 6)** | **Males w/ Fin**<br>**(n = 9)** | **p-value** |
| Body Mass (g) | 24.67 ± 0.36 | 23.53 ± 0.73 | 0.2562 |
| Bladder |  |  |  |
| Mass (% Body Mass) | 0.13 ± 0.01 | 0.12 ± 0.01 | 0.3701 |
| Volume (mm<sup>3</sup>/g Body Mass) | 33.71 ± 9.25 | 41.35 ± 6.61 | 0.5016 |
| **Prostate Mass (% Body Mass)** | **0.21 ± 0.01** | **0.14 ± 0.01** | **0.0009** |
| **Anterior Mass (% Body Mass)** | **0.11 ± 0.01** | **0.07 ± 0.00** | **0.0004** |
| **Ventral Mass (% Body Mass)** | **0.05 ± 0.01** | **0.04 ± 0.00** | **0.0467** |
| Dorsal Mass (% Body Mass) | 0.03 ± 0.00 | 0.03 ± 0.00 | 0.3465 |
| **Lateral Mass (% Body Mass)** | **0.02 ± 0.00** | **0.01 ± 0.00** | **0.0292** |
| **Seminal Vesicle Mass (% Body Mass)** | **0.72 ± 0.03** | **0.33 ± 0.03** | **<0.0001** |
| **Void Spot Assay** |  |  |  |
|  | **Males w/ Oil**<br>**(n = 7)** | **Males w/ Fin**<br>**(n = 9)** | **p-value** |
| Count | 23.20 ± 4.92 | 24.10 ± 10.10 | 0.1692 |
| Total Area (cm<sup>2</sup>) | 17.00 ± 4.33 | 18.40 ± 4.41 | 0.8209 |
| Percent area in center | 8.82 ± 7.54 | 12.10 ± 11.00 | 0.8845 |
| Percent area in corners | 48.90 ± 10.90 | 58.50 ± 13.30 | 0.3610 |
| Spots of a certain size (dia.) |  |  |  |
| 0 - 0.1 cm | 19.90 ± 4.36 | 21.00 ± 9.29 | 0.8658 |
| 0.1 - 0.25 cm | 0.89 ± 0.35 | 1.22 ± 0.64 | >0.9999 |
| 0.25 - 0.5 cm | 0.89 ± 0.31 | 0.22 ± 0.15 | 0.1312 |
| 0.5 - 1 cm | 0.11 ± 0.11 | 0.22 ± 0.22 | >0.9999 |
| 1 - 2 cm | 0.11 ± 0.11 | 0 ± 0 | >0.9999 |
| 2 - 3 cm | 0.11 ± 0.11 | 0.11 ± 0.11 | >0.9999 |
| 3 - 4 cm | 0.11 ± 0.11 | 0 ± 0 | >0.9999 |
| 4+ cm | 1.11 ± 0.31 | 1.33 ± 0.33 | 0.8013 |
| **Cystometry** |  |  |  |
|  | **Males w/ Oil**<br>**(n = 9)** | **Males w/ Fin**<br>**(n = 9)** | **p-value** |
| Void Duration | 0.63 ± 0.07 | 0.70 ± 0.13 | 0.8371 |
| **Intervoid Interval** | **3.35 ± 0.44** | **4.74 ± 0.42** | **0.0394** |
| Baseline Pressure | 3.96 ± 0.16 | 3.13 ± 0.51 | 0.1590 |
| Normalized Threshold Pressure | 3.64 ± 0.67 | 3.65 ± 0.56 | 0.9965 |
| Normalized Peak Void Pressure | 13.00 ± 0.99 | 15.90 ± 1.74 | 0.2011 |
| Non-voiding contractions | 1.21 ± 0.22 | 2.33 ± 0.53 | 0.0755<sup>Δ</sup> |
| **Voided Volume ( x 10<sup>-2</sup>)** | **4.24 ± 0.69** | **6.44 ± 0.47** | **0.0168** |
| Compliance ( x 10<sup>-2</sup>) | 1.24 ± 0.22 | 1.89 ± 0.31 | 0.1280 |
| Volume Flow Rate ( x 10<sup>-3</sup>) | 1.31 ± 0.27 | 1.87 ± 0.23 | 0.1377 |
| Mass Based Flow Rate | 22.80 ± 3.63 | 17.10 ± 1.74 | 0.1497 |
| Efficiency (%) | 108.00 ± 20.40 | 110.00 ± 6.85 | 0.8371 |

**Table 6.** Impact of Finasteride on Female Urinary Physiology.

| **Gross Assessment** | | Values are: Average ± SEM | |
| --- | --- | --- | --- |
|  | Female w/ Oil<br>(n = 7) | Female w/ Fin<br>(n = 5) | p-value |
| Body Mass (g) | 18.49 ± 0.42 | 17.50 ± 0.31 | 0.0953<sup>Δ</sup> |
| Bladder |  |  |  |
| Mass (% Body Mass) | 0.12 ± 0 | 0.12 ± 0.01 | >0.9999 |
| Volume (mm<sup>3</sup>/g Body Mass) | 17.82 ± 3.88 | 17.27 ± 3.24 | 0.9162 |
| **Void Spot Assay** |  |  |  |
|  | Female w/ Oil<br>(n = 7) | Female w/ Fin<br>(n = 5) | p-value |
| Count | 23.30 ± 5.73 | 12.20 ± 3.16 | 0.1337 |
| **Total Area (cm<sup>2</sup>)** | **22.20 ± 2.72** | **11.70 ± 1.51** | **0.0083** |
| Percent area in center | 0.65 ± 0.25 | 0.49 ± 0.33 | 0.1698 |
| Percent area in corners | 67.50 ± 12.60 | 73.70 ± 8.43 | 0.7023 |
| Spots of a certain size (dia.) |  |  |  |
| 0 - 0.1 cm | 17.90 ± 5.02 | 8.50 ± 2.68 | 0.1464 |
| 0.1 - 0.25 cm | 1.29 ± 0.71 | 0.50 ± 0.34 | 0.6166 |
| 0.25 - 0.5 cm | 0.14 ± 0.14 | 0.17 ± 0.17 | >0.9999 |
| 0.5 - 1 cm | 0.29 ± 0.18 | 0.33 ± 0.21 | >0.9999 |
| 1 - 2 cm | 0.29 ± 0.18 | 0.50 ± 0.34 | 0.7797 |
| 2 - 3 cm | 0.14 ± 0.14 | 0.83 ± 0.31 | 0.0862<sup>Δ</sup> |
| 3 - 4 cm | 0.71 ± 0.71 | 0 ± 0 | >0.9999 |
| **4+ cm** | **2.57 ± 0.297** | **1.33 ± 0.21** | **0.0087** |
| **Cystometry** |  |  |  |
|  | Female w/ Oil<br>(n = 7) | Female w/ Fin<br>(n = 5) | p-value |
| Void Duration | 0.51 ± 0.05 | 0.47 ± 0.03 | 0.5419 |
| Intervoid Interval | 2.90 ± 0.56 | 2.49 ± 0.30 | 0.5803 |
| Baseline Pressure | 5.70 ± 1.25 | 4.88 ± 0.83 | 0.6330 |
| Normalized Threshold Pressure | 11.3 ± 1.87 | 8.22 ± 0.75 | 0.2212 |
| Normalized Peak Void Pressure | 25.2 ± 1.32 | 21.70 ± 1.38 | 0.0965<sup>Δ</sup> |
| Non-voiding contractions | 3.63 ± 0.1.12 | 1.64 ± 0.47 | 0.3422 |
| Voided Volume ( x 10<sup>-2</sup>) | 4.10 ± 0.638 | 3.72 ± 0.42 | 0.6595 |
| Compliance ( x 10<sup>-2</sup>) | 0.29 ± 0.04 | 0.37 ± 0.06 | 0.2640 |
| Volume Flow Rate ( x 10<sup>-3</sup>) | 1.39 ± 0.20 | 1.37 ± 0.22 | 0.9335 |
| Mass Based Flow Rate | 17.90 ± 9.02 | 7.99 ± 1.10 | 0.4346 |
| Efficiency (%) | 111.00 ± 4.55 | 114.00 ± 6.44 | 0.7246 |

To address the impact of reducing prostate mass without surgery or hormone treatment, we used a genetic strategy to deplete prostatic luminal epithelial cells (Pbsn4^cre/+^; R26R^Dta/+^). Results are summarized in Figure 2-D and Table 7. C*re-*driven epithelial cell death reduces dorsal and lateral prostate mass without significantly changing mass of other prostate lobes or seminal vesicle. There were no statistical differences in VSA- or CMG-measured voiding function between Pbsn4^cre/+^; R26R^Dta/+^ mice and their genetic controls (Pbsn4^cre/+^; R26R^Td/+^).

**Table 7.** Impact of Genetic Prostatic Luminal Cell Ablation on Male Urinary Physiology.

| **Gross Assessment** | | Values are: Average ± SEM | |
| --- | --- | --- | --- |
|  | **Pbsn4<sup>cre/+</sup>; R26R<sup>Td/+</sup>**<br>(n = 8) | **Pbsn4<sup>cre/+</sup>; R26R<sup>Dta/+</sup>**<br>(n = 4) | **p-value** |
| Body Mass (g) | 25.58 ± 0.59 | 24.35 ± 0.80 | 0.1414 |
| Bladder |  |  |  |
| Mass (% Body Mass) | 0.15 ± 0.01 | 0.14 ± 0.00 | 0.5442 |
| Volume (mm<sup>3</sup>/g Body Mass) | 35.52 ± 6.16 | 36.98 ± 10.11 | 0.8992 |
| Prostate Mass (% Body Mass) | 0.22 ± 0.02 | 0.20 ± 0.01 | 0.3434 |
| Anterior Mass (% Body Mass) | 0.11 ± 0.00 | 0.12 ± 0.01 | 0.4629 |
| Ventral Mass (% Body Mass) | 0.05 ± 0.01 | 0.06 ± 0.00 | 0.6707 |
| **Dorsal Mass (% Body Mass)** | **0.05 ± 0.00** | **0.03 ± 0.00** | **0.0222** |
| **Lateral Mass (% Body Mass)** | **0.01 ± 0.00** | **0.00 ± 0.00** | **0.0242** |
| Seminal Vesicle Mass (% Body Mass) | 0.76 ± 0.04 | 0.72 ± 0.03 | 0.5431 |
| **Void Spot Assay** |  |  |  |
|  | **Pbsn4<sup>cre/+</sup>; R26R<sup>Td/+</sup>**<br>(n = 8) | **Pbsn4<sup>cre/+</sup>; R26R<sup>Dta/+</sup>**<br>(n = 4) | **p-value** |
| Count | 78.10 ± 30.09 | 73.00 ± 54.70 | 0.3677 |
| Total Area (cm<sup>2</sup>) | 36.60 ± 3.55 | 30.60 ± 6.96 | 0.4060 |
| Percent area in center | 5.47 ± 2.30 | 13.50 ± 13.30 | 0.2141 |
| Percent area in corners | 69.40 ± 6.49 | 56.00 ± 20.50 | 0.4413 |
| Spots of a certain size (dia.) |  |  |  |
| 0 - 0.1 cm | 67.30 ± 26.50 | 51.30 ± 36.10 | 0.3455 |
| 0.1 - 0.25 cm | 6.00 ± 4.02 | 14.00 ± 13.70 | 0.3939 |
| 0.25 - 0.5 cm | 1.25 ± 0.49 | 4.50 ± 4.17 | 0.8990 |
| 0.5 - 1 cm | 1.00 ± 0.38 | 1.25 ± 0.95 | >0.9999 |
| 1 - 2 cm | 0 ± 0 | 0 ± 0 | >0.9999 |
| 2 - 3 cm | 0 ± 0 | 0 ± 0 | >0.9999 |
| 3 - 4 cm | 0.13 ± 0.13 | 0.25 ± 0.25 | >0.9999 |
| 4+ cm | 2.50 ± 0.27 | 1.75 ± 0.25 | 0.1293 |
| **Cystometry** |  |  |  |
|  | **Pbsn4<sup>cre/+</sup>; R26R<sup>Td/+</sup>**<br>(n = 8) | **Pbsn4<sup>cre/+</sup>; R26R<sup>Dta/+</sup>**<br>(n = 4) | **p-value** |
| Void Duration | 0.44 ± 0.06 | 0.44 ± 0.07 | 0.9972 |
| Intervoid Interval | 5.32 ± 0.80 | 3.76 ± 0.87 | 0.2546 |
| Baseline Pressure | 2.44 ± 0.26 | 2.34 ± 0.62 | 0.7471 |
| Normalized Threshold Pressure | 5.68 ± 1.59 | 3.65 ± 0.77 | 0.4083 |
| Normalized Peak Void Pressure | 19.70 ± 3.51 | 16.70 ± 4.80 | 0.6277 |
| Non-voiding contractions | 6.60 ± 2.06 | 2.12 ± 0.57 | 0.0695<sup>Δ</sup> |
| Voided Volume (x 10<sup>-2</sup>) | 6.97 ± 1.09 | 6.14 ± 1.24 | 0.6720 |
| Compliance (x 10<sup>-2</sup>) | 1.81 ± 0.35 | 1.59 ± 0.452 | 0.7230 |
| Volume Flow Rate (x 10<sup>-3</sup>) | 3.69 ± 1.02 | 2.21 ± 0.214 | 0.3574 |
| Mass Based Flow Rate | 31.70 ± 2.49 | 19.90 ± 6.63 | 0.0672<sup>Δ</sup> |
| Efficiency (%) | 107.00 ± 8.21 | 114.00 ± 12.80 | 0.6029 |

The results of the three different methods of prostate mass reduction show the complex interplay of hormones and anatomical urinary tract variation that result in altered urinary function in male mice. We were surprised that the three methods of prostate reduction had inconsistent and minimal effects on urinary function. This is possibly due to variability in the localization and extent of prostate mass reduction among our three methods. In our companion paper, we explore this further through histologic evaluation of the prostate and urethra structure following each of our prostate mass manipulations. It is also noteworthy that we evaluated prostate mass reductions in our study; increases in prostate mass could have discrete impacts on urinary function.^37^

### 3.3 Exogenous Testosterone Supplementation Masculinizes Female Urinary Physiology

Because male castration feminized some of male urinary physiology, and because other methods of prostate mass reduction failed to recapitulate the resulting changes in male voiding function, we next tested if circulating testosterone underlies sex differences in mouse urinary function. We supplemented female mice with exogenous testosterone. We assessed the supplemented mice for directional changes in VSA/CMG parameters consistent with “masculinization” of urinary physiology (i.e. the parameters changed in the same direction as males). Female mice were divided into two groups: control or with silastic testosterone capsule implants (Female w/ T). Capsules were implanted at 6 weeks of age and assessment took place at 9 weeks of age. Results are summarized in Figure 4-B and Table 8.

**Figure 4.**
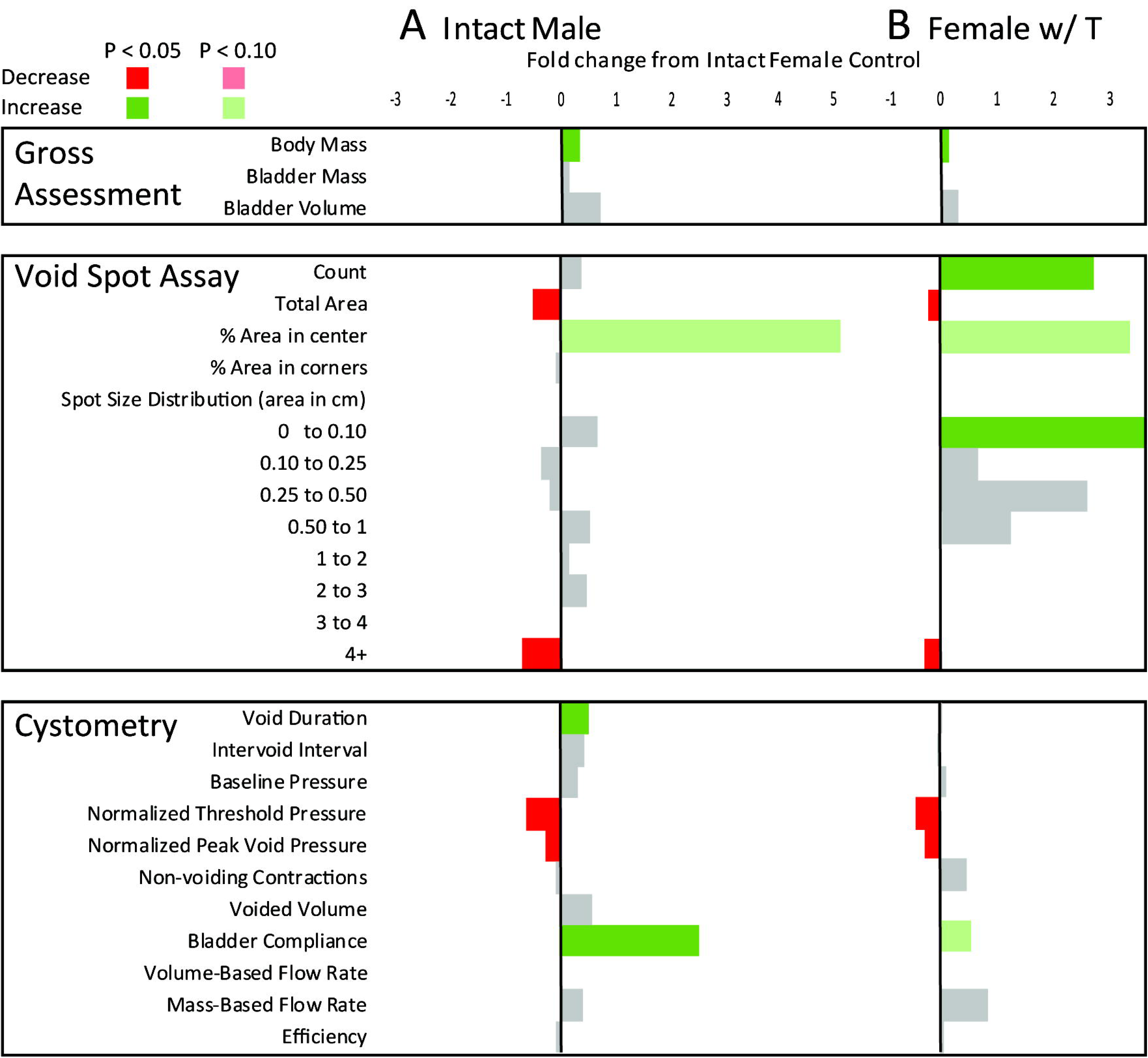
Summary statistics for anatomical and physiological differences across experimental groups evaluated in this study including intact male and female mice, and female mice implanted with 1 cm silastic capsules containing crystalline testosterone (Female w/ T). All results are mean fold differences relative to intact female control. Differences that are significantly higher than intact females are shown in green and those that are lower than intact females are in red. Endpoints that do not significantly differ from intact females are in gray. The numerical fold difference relative to intact females is indicated by the bar size according to the legend at the top of the figure. Male urinary physiology is different from female at baseline. The female voiding parameters affected by exogenous testosterone include many of the same parameters that distinguish male from female voiding function.

**Table 8.**
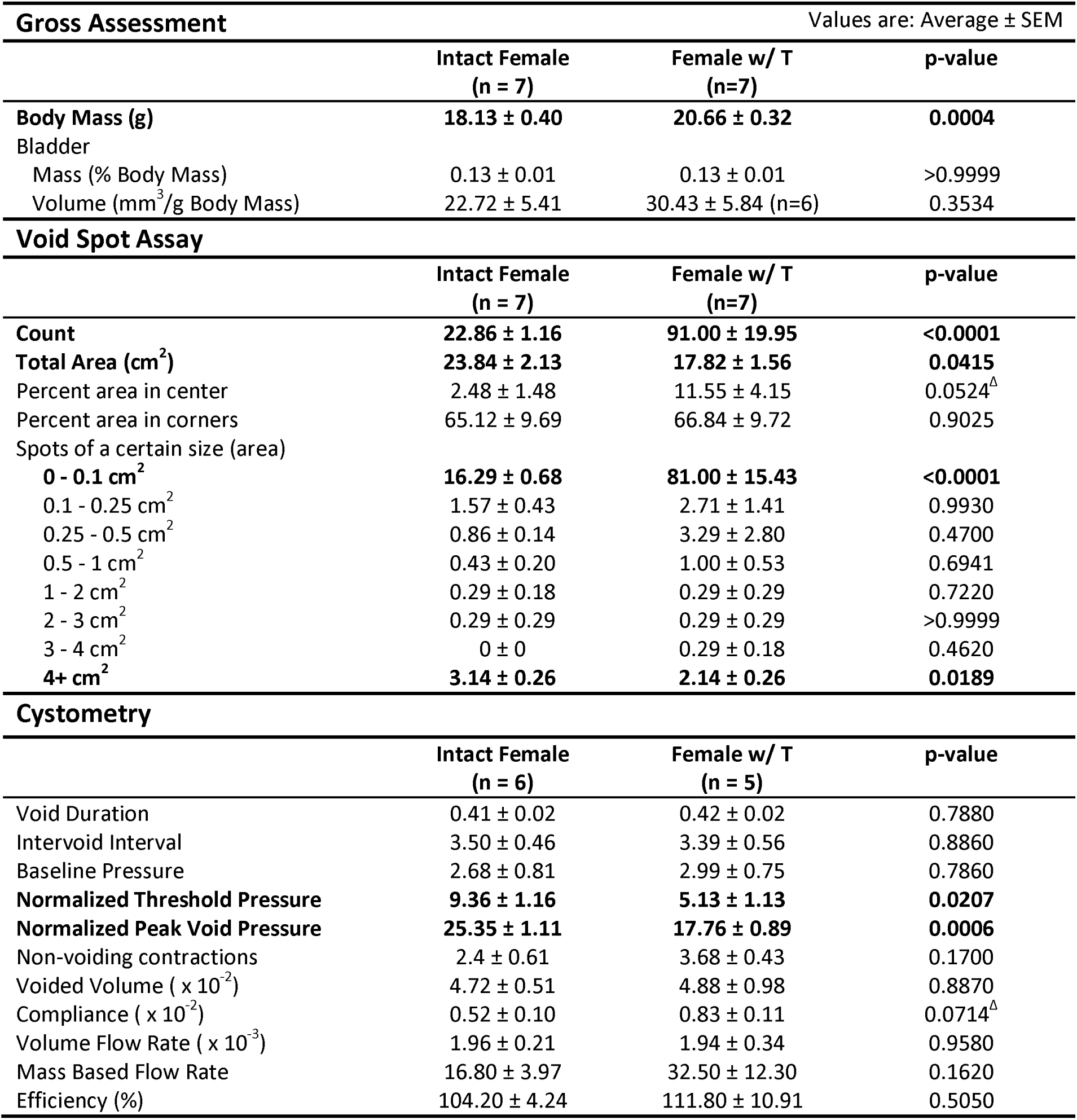
Impact of Testosterone on Female Urinary Physiology.

Females w/ T have a significantly greater body mass but similar bladder mass and volume compared to female controls. A previous study reported that testosterone does not change female mouse body weight but increases bladder weight.^35^ However, this study used mice of a different age, strain, environment, and method of testosterone delivery.

Testosterone supplementation caused several changes in female voiding function measured by VSA. Similar to intact males, Female w/ T mice deposit significantly less urine during a four hour monitoring period, void fewer spots greater than 4 cm^2^, and void a larger percentage in the center of the paper than female controls. Also similar to males, Female w/ T mice have a significantly lower threshold pressure, a lower peak void pressure, and higher bladder compliance than control females. Females w/ T, unlike males, have more void spots -- specifically more 0 to 0.1cm^2^ spots than female controls.

Female voiding parameters affected by exogenous testosterone include many of the same parameters that distinguish male from female voiding function. Thus, we conclude that even though female lower urinary tract anatomy differs from that of males, exogenous testosterone “masculinizes” female voiding patterns. The clear relationship between testosterone and sex differences in urinary function was a surprising finding of this study. Androgen receptor expression and activity is well described in the reproductive tract, but its expression and activity in the urinary tract isn’t well characterized. A previous report documented androgen receptor expression in occasional stromal cells of the bladder and weak staining in the kidney.^38^ Another found that pelvic ganglia contain androgen sensitive autonomic nerves.^39^ This finding raises the possibility that androgens can masculinize autonomic signaling in the female lower urinary tract to drive urinary voiding patterns similar to intact male mice. Further studies to compare androgen receptor expression of the entire male and female lower urinary tract are needed to better elucidate the influence of androgens on autonomic signaling in the bladder, prostate and urethra.

A VSA feature not accounted for by “masculinization” of urinary function was the increase in spot count, specifically small spots (0-0.1 cm^2^). We implanted testosterone capsules into 8-week-old male mice (Male w/ T) and assessed voiding one week later (Table 9). Males w/ T trend toward an increase in total void spot count, p=0.0760, and small (0-0.1 cm^2^) spot count, p=0.0696, compared to male controls, making increased voiding frequency a common feature of exogenous testosterone supplementation in females and intact males (overall p<0.0001 and p=0.0760, small p<0.0001 and p=0.0696).

**Table 9.**
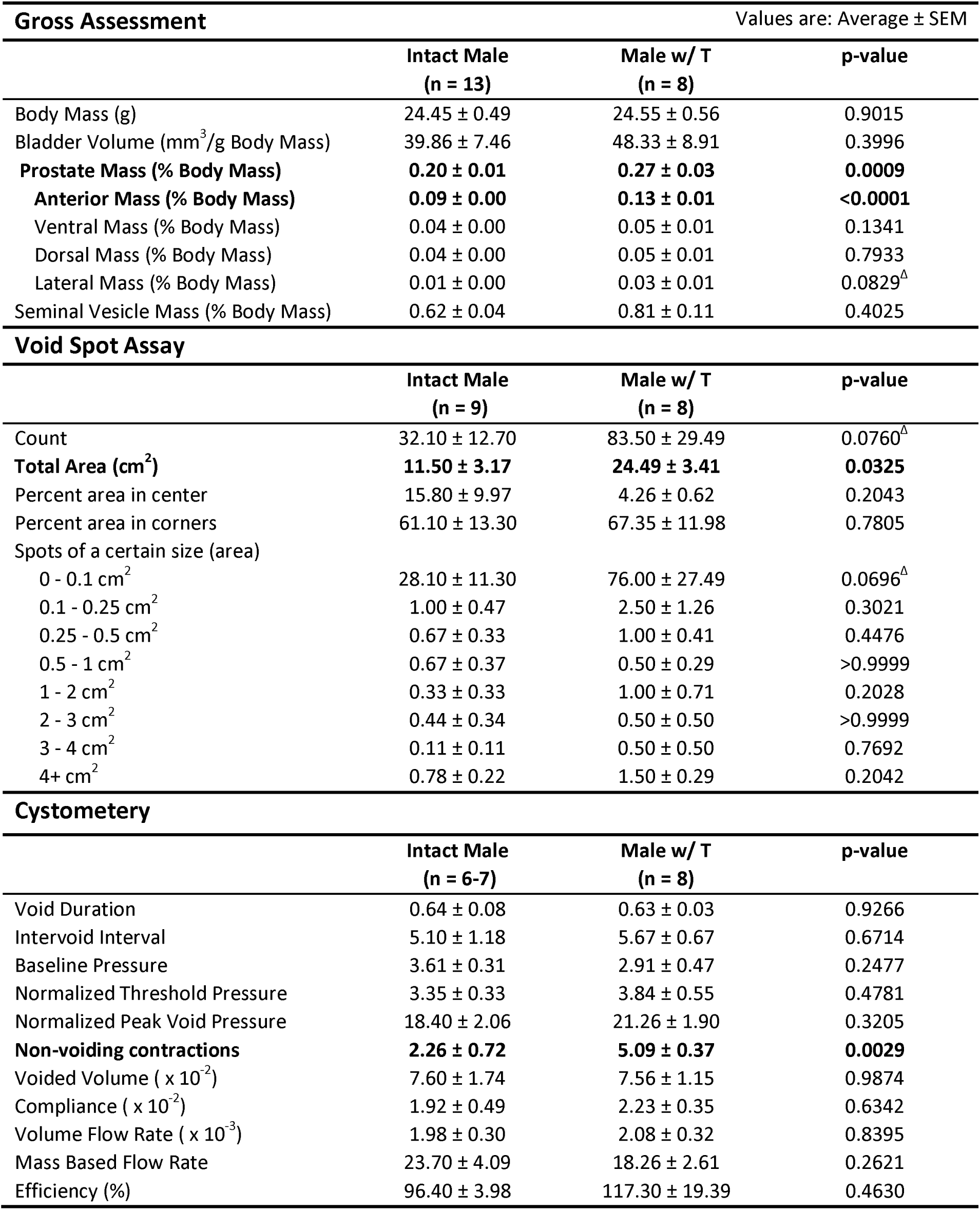
Impact of Testosterone on Intact Male Urinary Physiology.

| **Gross Assessment** | | Values are: Average ± SEM | |
| --- | --- | --- | --- |
|  | **Intact Male**<br>**(n = 13)** | **Male w/ T**<br>**(n = 8)** | **p-value** |
| Body Mass (g) | 24.45 ± 0.49 | 24.55 ± 0.56 | 0.9015 |
| Bladder Volume (mm<sup>3</sup>/g Body Mass) | 39.86 ± 7.46 | 48.33 ± 8.91 | 0.3996 |
| **Prostate Mass (% Body Mass)** | **0.20 ± 0.01** | **0.27 ± 0.03** | **0.0009** |
| **Anterior Mass (% Body Mass)** | **0.09 ± 0.00** | **0.13 ± 0.01** | **<0.0001** |
| Ventral Mass (% Body Mass) | 0.04 ± 0.00 | 0.05 ± 0.01 | 0.1341 |
| Dorsal Mass (% Body Mass) | 0.04 ± 0.00 | 0.05 ± 0.01 | 0.7933 |
| Lateral Mass (% Body Mass) | 0.01 ± 0.00 | 0.03 ± 0.01 | 0.0829<sup>Δ</sup> |
| Seminal Vesicle Mass (% Body Mass) | 0.62 ± 0.04 | 0.81 ± 0.11 | 0.4025 |
| **Void Spot Assay** |  |  |  |
|  | **Intact Male**<br>**(n = 9)** | **Male w/ T**<br>**(n = 8)** | **p-value** |
| Count | 32.10 ± 12.70 | 83.50 ± 29.49 | 0.0760<sup>Δ</sup> |
| **Total Area (cm<sup>2</sup>)** | **11.50 ± 3.17** | **24.49 ± 3.41** | **0.0325** |
| Percent area in center | 15.80 ± 9.97 | 4.26 ± 0.62 | 0.2043 |
| Percent area in corners | 61.10 ± 13.30 | 67.35 ± 11.98 | 0.7805 |
| Spots of a certain size (area) |  |  |  |
| 0 - 0.1 cm<sup>2</sup> | 28.10 ± 11.30 | 76.00 ± 27.49 | 0.0696<sup>Δ</sup> |
| 0.1 - 0.25 cm<sup>2</sup> | 1.00 ± 0.47 | 2.50 ± 1.26 | 0.3021 |
| 0.25 - 0.5 cm<sup>2</sup> | 0.67 ± 0.33 | 1.00 ± 0.41 | 0.4476 |
| 0.5 - 1 cm<sup>2</sup> | 0.67 ± 0.37 | 0.50 ± 0.29 | >0.9999 |
| 1 - 2 cm<sup>2</sup> | 0.33 ± 0.33 | 1.00 ± 0.71 | 0.2028 |
| 2 - 3 cm<sup>2</sup> | 0.44 ± 0.34 | 0.50 ± 0.50 | >0.9999 |
| 3 - 4 cm<sup>2</sup> | 0.11 ± 0.11 | 0.50 ± 0.50 | 0.7692 |
| 4+ cm<sup>2</sup> | 0.78 ± 0.22 | 1.50 ± 0.29 | 0.2042 |
| **Cystometry** |  |  |  |
|  | **Intact Male**<br>**(n = 6-7)** | **Male w/ T**<br>**(n = 8)** | **p-value** |
| Void Duration | 0.64 ± 0.08 | 0.63 ± 0.03 | 0.9266 |
| Intervoid Interval | 5.10 ± 1.18 | 5.67 ± 0.67 | 0.6714 |
| Baseline Pressure | 3.61 ± 0.31 | 2.91 ± 0.47 | 0.2477 |
| Normalized Threshold Pressure | 3.35 ± 0.33 | 3.84 ± 0.55 | 0.4781 |
| Normalized Peak Void Pressure | 18.40 ± 2.06 | 21.26 ± 1.90 | 0.3205 |
| **Non-voiding contractions** | **2.26 ± 0.72** | **5.09 ± 0.37** | **0.0029** |
| Voided Volume ( x 10<sup>-2</sup>) | 7.60 ± 1.74 | 7.56 ± 1.15 | 0.9874 |
| Compliance ( x 10<sup>-2</sup>) | 1.92 ± 0.49 | 2.23 ± 0.35 | 0.6342 |
| Volume Flow Rate ( x 10<sup>-3</sup>) | 1.98 ± 0.30 | 2.08 ± 0.32 | 0.8395 |
| Mass Based Flow Rate | 23.70 ± 4.09 | 18.26 ± 2.61 | 0.2621 |
| Efficiency (%) | 96.40 ± 3.98 | 117.30 ± 19.39 | 0.4630 |

To determine whether restoration of physiologic testosterone induces voiding frequency in castrate mice, we implanted castrate males with testosterone capsules (Castrate w/ T) at eight weeks of age and assessed at nine weeks of age (Table 10). Spot count did not increase when castrate mice were supplemented with testosterone. Therefore, testosterone increases voiding frequency when increased to supra-physiologic concentrations, but not when depleted and then returned to physiologic concentrations.

**Table 10.** Impact of Testosterone on Castrate Male Urinary Physiology.

| **Gross Assessment** | Values are: Average ± SEM | | |
| --- | --- | --- | --- |
|  | **Castrate Male**<br>**(n = 14)** | **Castrate w/ T**<br>**(n=8)** | **p-value** |
| **Body Mass (g)** | **21.51 ± 0.41** | **23.65 ± 0.69** | **0.0023** |
| **Bladder** |  |  |  |
| Volume (mm<sup>3</sup>/g Body Mass) | 18.04 ± 3.13 | 33.57 ± 6.89 | 0.0192 |
| Prostate Mass (% Body Mass) | 0.03 ± 0.00 | 0.21 ± 0.02 | <0.0001 |
| Anterior Mass (% Body Mass) | 0.02 ± 0.00 | 0.10 ± 0.01 | <0.0001 |
| Ventral Mass (% Body Mass) | 0.01 ± 0.00 | 0.05 ± 0.01 | <0.0001 |
| Dorsal Mass (% Body Mass) | 0.01 ± 0.00 | 0.04 ± 0.00 | <0.0001 |
| Lateral Mass (% Body Mass) | 0.00 ± 0.00 | 0.02 ± 0.00 | 0.0009 |
| Seminal Vesicle Mass (% Body Mass) | 0.04 ± 0.00 | 0.35 ± 0.07 | <0.0001 |
| **Void Spot Assay** |  |  |  |
|  | **Castrate Male**<br>**(n = 11)** | **Castrate w/ T**<br>**(n = 4)** | **p-value** |
| Count | 42.00 ± 6.10 | 50.30 ± 14.90 | 0.5462 |
| Total Area (cm<sup>2</sup>) | 20.50 ± 2.31 | 22.60 ± 8.62 | 0.7419 |
| Percent area in center | 2.34 ± 0.666 | 7.35 ± 5.93 | 0.8645 |
| Percent area in corners | 77.40 ± 3.96 | 72.80 ± 15.30 | 0.6846 |
| Spots of a certain size (area) |  |  |  |
| 0 - 0.1 cm<sup>2</sup> | 38.10 ± 5.84 | 44.00 ± 13.20 | 0.6410 |
| 0.1 - 0.25 cm<sup>2</sup> | 1.36 ± 0.43 | 2.00 ± 0.82 | 0.5509 |
| 0.25 - 0.5 cm<sup>2</sup> | 0.64 ± 0.15 | 1.00 ± 1.00 | 0.5692 |
| 0.5 - 1 cm<sup>2</sup> | 0.46 ± 0.21 | 0 ± 0 | 0.3956 |
| 1 - 2 cm<sup>2</sup> | 0.18 ± 0.12 | 0.25 ± 0.25 | >0.9999 |
| 2 - 3 cm<sup>2</sup> | 0 ± 0 | 0.50 ± 0.50 | 0.2667 |
| 3 - 4 cm<sup>2</sup> | 0 ± 0 | 0 ± 0 | >0.9999 |
| 4+ cm<sup>2</sup> | 1.27 ± 0.14 | 2.50 ± 0.87 | 0.1692 |
| **Cystometry** |  |  |  |
|  | **Castrate Male**<br>**(n = 7-8)** | **Castrate w/ T**<br>**(n = 8)** | **p-value** |
| Void Duration | 0.42 ± 0.03 | 0.51 ± 0.04 | 0.1049 |
| Intervoid Interval | 2.77 ± 0.12 | 3.00 ± 0.45 | 0.6444 |
| Baseline Pressure | 2.21 ± 0.26 | 2.92 ± 0.61 | 0.3111 |
| Normalized Threshold Pressure | 4.75 ± 0.73 | 3.32 ± 0.58 | 0.1437 |
| Normalized Peak Void Pressure | 17.20 ± 0.75 | 16.00 ± 1.70 | 0.5400 |
| **Non-voiding contractions** | **1.46 ± 0.25** | **3.58 ± 0.56** | **0.0039** |
| Voided Volume ( x 10<sup>-2</sup>) | 4.05 ± 0.11 | 4.60 ± 0.64 | 0.4184 |
| Compliance ( x 10<sup>-2</sup>) | 0.89 ± 0.20 | 1.34 ± 0.31 | 0.1840 |
| Volume Flow Rate ( x 10<sup>-3</sup>) | 1.69 ± 0.16 | 1.53 ± 0.18 | 0.5159 |
| Mass Based Flow Rate | 23.40 ± 3.17 | 20.60 ± 4.57 | 0.6126 |
| Efficiency (%) | 96.80 ± 3.31 | 90.74 ± 5.81 | 0.3993 |

A “frequent voider” pattern, as detected by VSA (>100 urine spots deposited on a filter paper in a four hour monitoring period), was previously noted in approximately 10% of 9-week-old C57BL/6J mice.^29^ In our study, 14.29% of testosterone treated females and 25% of testosterone treated males were frequent voiders, but no control females and just 11% of control males were frequent voiders. These results further support the notion that supra-physiologic testosterone increases voiding frequency and are consistent with increased voiding frequency in rats and dogs supplemented with testosterone.^40^

## 4. CONCLUSIONS

We used surgical, pharmacological, and genetic approaches to reduce mouse prostate mass, and also exposed C57BL/6J mice to exogenous testosterone to determine the influence of the prostate and testosterone on voiding function. We characterized male and intact female urinary phenotypes using contemporary methodologies and highlighted unique sex differences in urinary voiding phenotype.

Urologic researchers and practitioners are beginning to appreciate that not all male LUTS arise from urethral occlusion by an enlarged prostate and that additional factors (prostatic urethra collagen accumulation,^12^ prostatic inflammation,^13^ and prostate smooth muscle contraction^14, 15^) also drive LUTS. Mice are instrumental in studying these alternative mechanisms. It is possible, for example, to induce prostatic inflammation and drive urinary frequency and pelvic pain in mice modeling LUTS symptomology.^22, 41, 42^ It is also possible to knock-out or overexpress genes and cell types to validate mechanisms arising from clinical studies and to use mice for pre-clinical safety and efficacy trials of novel therapeutics. Mice fuel a creative cycle of translational research resulting in specific targeted therapies for men. In in order to take full advantage of our models, further baseline studies are needed to determine baseline molecular and cellular contributions to voiding function in our mouse models.

## ACKNOWLEDGEMENTS

Funded by National Institutes of Health grants: U54 DK104310, Summer Program In Undergrduate Urologic Research (U54 DK104310S1), R01ES001332, R01DK099328, F31ES028594, TL1TR002375, and University of Wisconsin-Madison, School of Veterinary Medicine. The content is solely the responsibility of the authors and does not necessarily represent the official views of the National Institutes of Health.

